# Projections from subfornical organ to infralimbic cortex modulate carbon dioxide associated fear

**DOI:** 10.64898/2026.05.15.725536

**Authors:** Katherine M.J. McMurray, Andrew Winter, Rebecca Ahlbrand, Sachi Shukla, Aravind Kalathill, Andrew Gaulden, Benjamin Packard, Allan-Hermann Pool, Steve Davidson, Jayme R. McReynolds, James P. Herman, Renu Sah

## Abstract

Most of our mechanistic understanding of threat responding and defensive fear behaviors is based on exposure to aversive stimuli in the environment. However, unpleasant, within-the body interoceptive signals can also regulate threat and emotion although underlying cell-circuit mechanisms are not well understood. Abnormal interoceptive sensitivity is associated with fear-associated psychiatric conditions such as panic disorder and PTSD. The ventromedial infralimbic (IL) subdivision of the prefrontal cortex plays a key role in threat appraisal and fear, however, IL engagement in interoceptive threat response and contributory afferent mechanisms are not known. Here, using an interoceptive clinical panicogen, carbon dioxide (CO_2_) inhalation, we report IL-mediated regulation of fear in mice via afferents from the subfornical organ (SFO), a key viscero-humoral circumventricular organ lacking a traditional blood brain barrier. Chemogenetic inhibition of SFO-to-IL (but not SFO-to-BNST) projections regulated defensive behaviors during CO_2_ inhalation and associative contextual fear. Notably, the SFO-IL circuit also modulated delayed CO_2_ effects on contextual fear conditioning-extinction, but not startle, neuroendocrine response or motivated behaviors. We also established more specifically that SFO angiotensin II receptor type-1 (AT-1R)^+ve^ neuronal afferents to the IL regulate CO_2_-associated fear and long-term deficits in contextual fear extinction. CO_2_ inhalation reduced neuronal activation within the IL and optogenetic activation of SFO neurons activated inhibitory parvalbumin (PV) (but not somatostatin (SST)) interneurons in the IL. Collectively, these data reveal that aversive interoceptive signals can be directly conveyed to the IL via the SFO, a sensory hub for systemic perturbations, to regulate spontaneous and long-term fear. Our findings provide important mechanistic insights into fear-associated disorders with abnormal interoceptive threat sensitivity such as panic disorder and PTSD.

## Introduction

An organism’s ability to respond to stimuli that signal threat is crucial for survival. Most of our mechanistic understanding of threat responding and defensive fear behaviors is based on aversive stimuli in the environment, that are exteroceptive in nature^1,2^. However, unpleasant, within-the body sensations also present as threatful “interoceptive” signals to the brain. Abnormal interoceptive signaling is a hallmark of many fear-associated disorders such as panic disorder ^3^ and posttraumatic stress disorder (PTSD) ^4^ . Individuals with panic and PTSD often experience high emotional reactivity to threatful interoceptive triggers such as CO_2_ inhalation^3,5–7^ and CO_2_ sensitivity is associated with emergence of PTSD symptoms to war-zone trauma^8^ . Currently, mechanisms underlying the translation of interoceptive threat exposure into spontaneous and conditioned emotional behaviors are not well understood at the cell-circuit level.

The infralimbic (IL) subdivision of the medial prefrontal cortex (mPFC), considered as homologous to the ventromedial area 25 of the prefrontal cortex in humans^9^ is a key regulator of threat appraisal and associated defensive behaviors ^10^. Previous work has highlighted an important role of the IL in processing of emotional and homeostatic cues ^11^, detection of arousal ^12^ and signaling safety ^13^. IL-mediated regulation of environmental threat-associated fear is well understood ^14–16^, however, it is not known whether the IL is engaged by threatful interoceptive triggers such as CO_2_ inhalation that are relevant to the neurobiology of PD and PTSD. Furthermore, IL modulation of defensive behaviors via “top-down” regulation is well understood ^17,18^, however, afferent inputs and circuitry that can relay interoceptive threat signals to the IL are not known.

Circumventricular organs (CVOs) are brain nodes located in proximity to ventricles and lack a typical blood brain barrier (BBB)^19^. Among CVOs, sensory CVOs are unique in that they not only have access to systemic compartments but also possess strategic neural connections to effector target sites regulating emotional behaviors^20^. The subfornical organ (SFO), a sensory CVO located near the lateral ventricle regulates motivated behaviors that enable survival to homeostatic threats signaling dehydration, hypertonicity and hunger ^21–23^. Previously, we identified the SFO as a chemosensory site for detecting CO_2_-associated hypercapnia promoting fear and panic behavior^24^. Thus, in addition to regulating motivated behaviors to metabolic cues, the SFO can regulate emotional behaviors to fear-evoking interoceptive cues. Direct SFO projections to threat regulatory nodes such as the IL and BNST have been reported^20^ and recent studies show their role in inflammation-induced anxiety^25^ and fear^26^. SFO neuronal projections that regulate CO_2_-associated fear have not been identified.

With this background, we sought to examine projections from the SFO to the IL and BNST in regulating spontaneous and conditioned fear to CO_2_ inhalation. Given CO_2_ associations with later symptoms of PTSD^8^, we investigated delayed CO_2_ effects on fear conditioning and extinction as reported previously^27^. We also hypothesized a role of angiotensin II receptor type 1, AT-1R ^+ve^ SFO projections in CO_2_ behaviors, based on previous data showing AT-1R association with panic^28^, PTSD^29,30^ and CO_2_ inhalation ^31^. Our data revealed that SFO-IL (but not SFO-BNST) projections regulate CO_2_ associated spontaneous and conditioned fear as well as delayed effects on fear conditioning-extinction; while startle, neuroendocrine stress response or thirst-associated motivated behavior are not impacted. We further identify SFO AT1R^+ve^ IL projections in regulating CO_2_ behavioral effects.

## Materials and Methods

### Animals

Studies were performed using 8-week-old male BALB/c mice (Envigo, Indianapolis, IN) as reported in previous CO_2_-evoked behavior studies by our group ^24,27,32^ or angiotensin II type 1-receptor (AT1R) cre mice which express cre-recombinase under the AT1R promotor (is B6(C3)-*Agtr1a^tm1.1(cre)Ekrs^*/Jax Stock No: 030553) as indicated. Detection of AT-1R expression in the SFO was conducted in AT1aR-tdTomato reporter mice (gift from Dr. Eric Krause, Georgia State University, Atlanta, GA). Mice were group housed in a climate-controlled vivarium (temperature 23 ± 4 ^0^C, humidity 30 ± 6%) on a 14hr/10hr light/dark cycle. All study protocols were approved by the Institutional Animal Care and Use Committee of the University of Cincinnati and performed in a facility accredited by the Association for Assessment and Accreditation of Laboratory Animal Care.

### Surgeries

Mice were anesthetized using isoflurane and received viral infusions as follows. For experiments targeting SFO to IL projections, mice received bilateral infusions of a retrogradely transported cre-expressing virus (300 nl; AAVrg-pmSyn1-EBFP-Cre, RRID:Addgene_51507) targeting the IL (AP: 1.7mm, DV: -2.5, ML ± 0.3mm) and either infusion of a cre-dependent inhibitory DREADD virus (300 nl; AAV2-hSyn-DIO-hM4D(Gi)-mCherry, RRID:Addgene_44362) or cre-dependent mCherry control virus (300 nl; AAV2-hSyn-DIO-mCherry, RRID:Addgene_50459) targeting the SFO (8° angle, AP: -0.2 mm, DV: -3.1 mm, ML +0.5 mm). For experiments targeting AT1R expressing neurons that project from SFO to IL, AT1R cre mice received infusions of a cre-dependent recombinase FlpO (AAV5-EF1a-DIO-FLPo-WPRE-hGHpA; RRID: Addgene_87306) into the SFO. They also received bilateral infusions of a retrogradely transported flp-dependent inhibitory DREADD virus (AAVrg-hSyn-fDIO-hM4D(Gi)-mCherry-WPREpA;RRID:Addgene_154867) or control virus expressing flp-dependent mCherry (AAVrg-Ef1a-fDIO mCherry, RRID:Addgene_114471) targeted to the IL.

For experiments targeting SFO to BNST projections, mice received bilateral infusions of a retrogradely transported cre-expressing virus (AAVrg.hSyn.HI.eGFP-Cre.WPRE.SV40, RRID:Addgene_105540) targeting the BNST (AP: 0.3 mm, DV: -4.6 mm, ML ± 1.0 mm) ^22^and either infusion of the same cre-dependent inhibitory DREADD virus (300 nl) or cre-dependent mCherry control virus (300 nl;) to the SFO.

Mice were allowed to recover for 4 weeks before experimental testing began. Mice received a single injection of vehicle (0.133% DMSO in saline) or clozapine-n-oxide (CNO; 3mg/kg in vehicle; NIMH C-929) 30 minutes prior to CO_2_ inhalation, water consumption, or restraint stress (see below for details).

For experiments investigating the effects of SFO activation on neuronal activity within IL we targeted SFO projection neurons that are primarily excitatory CAMK-II^+ve^ ^33^. Mice received 50nl infusions of AAV5-CaMKIIα-hChR2(H134R)-mCherry (RRID:Addgene_26975) or AAV5-CaMKIIα-EGFP (RRID:Addgene_50469) virus into the SFO (see coordinates above). Animals were allowed to recover for two weeks and then subjected to surgeries for implantation of optogenetic probes (Plexon, 94091). Experiments began following 2 weeks of recovery.

For experiments using *in vivo* fiber photometry to record IL Ca^2+^ (as a proxy for neuronal activity) during CO_2_ inhalation, mice received unilateral infusions of 300 nl AAV1.CamKII.GCaMP6f.WPRE.SV40 (RRID:Addgene_100834) into IL (see coordinates above). A fiberoptic cannula (400 µm, NA0.48, 4mm; Doric Lenses, Quebec, Canada) was then implanted just dorsal to the viral infusion site AP: 1.7mm, DV: -2.4, ML ± 0.3mm. Mice were allowed to recover for 4 weeks before testing

### Behavioral manipulations

We used a paradigm developed by our lab ^27,34^ for assessment of passive and active defensive behaviors (freezing and rearing) to interoceptive threat, CO_2_, followed by delayed effects of prior CO_2_ inhalation on defensive responding to discrete exteroceptive cues (acoustic startle, foot shock contextual fear conditioning-extinction). We confirmed significant effects of CO_2_ on freezing during inhalation, context re-exposure and on fear conditioning and extinction one week later (Suppl Fig 2) as in our previous studies^27,34^. As the primary objective of the study was to investigate behaviors associated with interoceptive threat, all behavioral experiments were confined to assessing the effect of manipulations on CO_2_ (and not air)-evoked behaviors. As noted above, mice received a single injection of VEH/CNO 30 min prior to a single exposure to 5% CO_2_ inhalation. To assess delayed effects of CO_2_, foot shock contextual fear conditioning-extinction-reinstatement testing was conducted one week later. Effects of SFO to IL or SFO to BNST inhibition on post dehydration water consumption were assessed after one week. After 5 days restraint stress induced hypothalamic pituitary adrenal (HPA) axis response was measured. Mice received CNO injections 30 minutes prior to water access or restraint stress exposure

### CO_2_-inhalation paradigm

After 4 weeks post-surgery mice were subjected to CO_2_ inhalation (Fig 1 schematic) as described in previous studies by our group and others ^24,27,34,35^ using a dual vertical Plexiglas chamber (25.5cm x 29cm x 28cm per chamber). 5% (BALB/c mice) or 10% (AT1R-cre C57/Bl6 mice) CO_2_ (in 21% O_2_, balanced with N_2_, Wright Brothers Inc., Cincinnati, OH) was infused in the upper chamber while the mice were placed in the lower compartment to avoid direct blowing of the gas, which is aversive to rodents. A flow meter with a steady infusion rate of 10 L/min was used for all animals and the concentration of CO_2_ within the lower chamber was verified (5.0 ± 0.5%) by the CARBOCAP® GM70 carbon dioxide meter (GMP221 probe with accuracy specification +/- 0.5%) (Vaisala, Helsinki, Finland). The choice of 5% or 10% CO_2_ exposure was based on strain-dependent sensitivity to CO_2_ concentration (5% for BALB/c versus 10% for C57/Bl6 mice) as reported in our previous work^24,34,36^.

**Figure 1.**
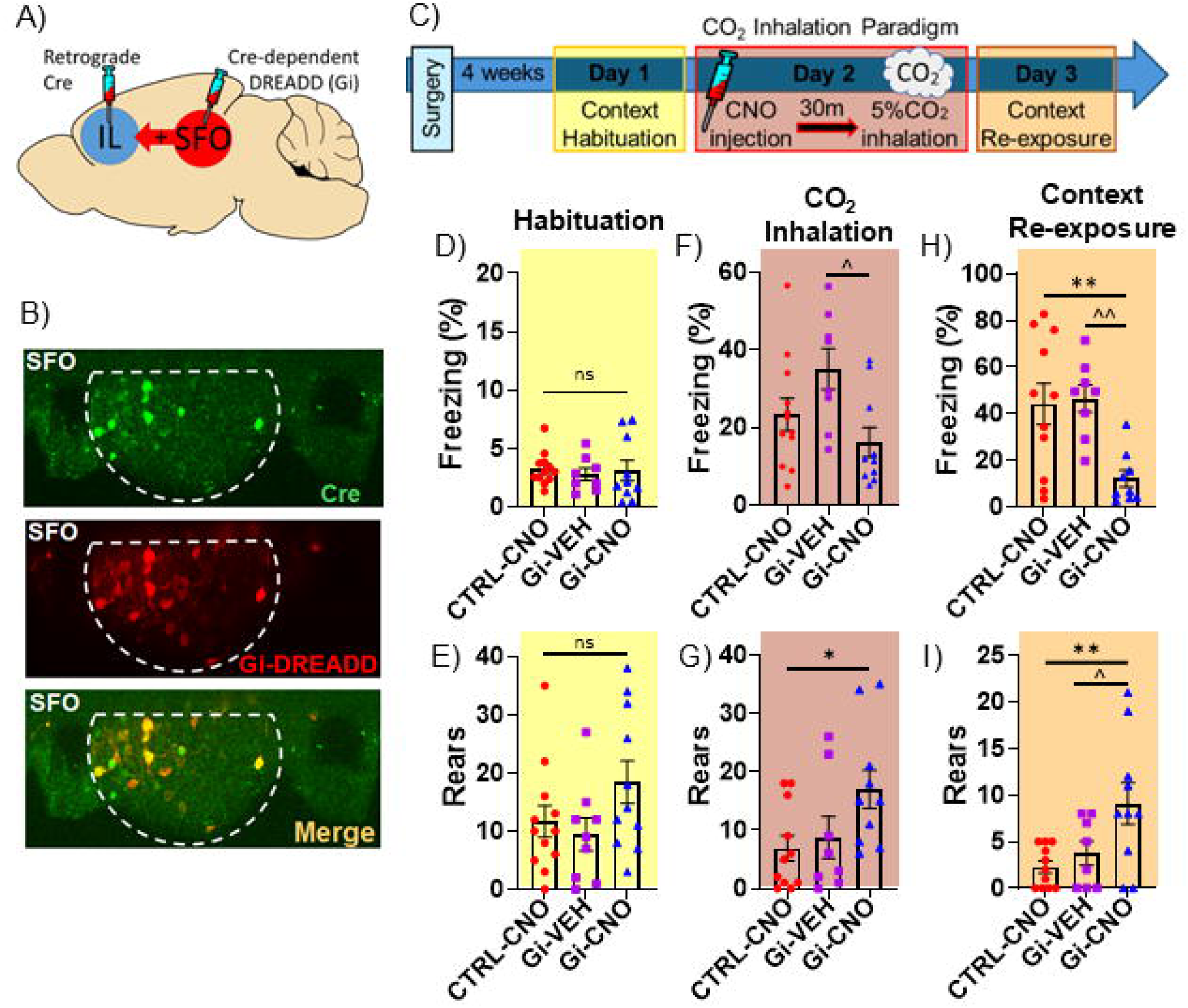
SFO to IL projection regulates defensive behaviors to interoceptive threat, CO_2_ inhalation. **(A)** Intersectional chemogenetics strategy to target the SFO-IL circuit using a Cre-expressing retrogradely transported viral construct - AAV2.hSyn.HI.eGFP-Cre.WPRE.SV40 into the IL and AAV2-hSyn-DIO-hM3D(Gi)-mCherry virus in the SFO to enable Cre-dependent expression of DREADDs in SFO neurons projecting to the IL. (**B**) Images from the SFO show the expression of Cre-eGFP (green) in Cre expressing SFO to IL neurons, Cre-dependent Gi-DREADD-mCherry or control mCherry virus expressing red fluorescent protein soma (red) and colocalized cells in merged image. **(C)** Schematic for experimental setup. Following surgeries mice were allowed to recover for 4 weeks before undergoing the CO_2_ inhalation paradigm (see text for details). Mice underwent context habituation on day 1. On day 2 a single CNO or vehicle injection was administered 30 min prior to CO_2_ inhalation (10 min) and were then re-exposed to inhalation context the next day in the absence of CO_2_. For habituation, no significant group difference was observed in freezing (**D**) or rearing (**E**). During 5% CO_2_ inhalation Gi-CNO mice elicited a reduction in freezing (**F**) and increase in rearing (**G**) compared to Gi-veh and Control virus (CTRL-CNO) groups. Following Day 3 context re-exposure for conditioned behaviors CNO-treated Gi-DREADD mice showed significantly lower freezing (**H**) and increased rearing (I) compared to control virus and vehicle groups. Data are expressed as mean ± sem (n= 8-12 mice/group); * p<0.05 **p<0.01 between CTRL-CNO and Gi-CNO; ^ p<0.05 ^^ p<0.01 between Gi-VEH and Gi-CNO, ns=not significant.

Mice were acclimated to a CO_2_ chamber (Day 0) during which baseline passive (freezing) and active (rearing) were assessed. The following day (Day 1), mice were returned to the chamber and exposed to CO_2_ inhalation for 10 min. The day after CO_2_ exposure (Day 2), animals were returned to the chamber for 5 min in the absence of CO_2_.for context conditioned behaviors. Mice were video recorded for later analysis. Freezing and rearing behaviors on Day 0-2 were analyzed Freezing (complete lack of movement except for respiration) was scored using the FreezeScan software (CleverSys Inc.) and rearing (standing on hind legs with or without foreleg contact on wall) was scored by a trained observer blinded to experimental condition. CO_2_-evoked freezing behavior appears to be consistent in rodents and likely represents a fear-associated defensive behavior. Rearing frequency has been reported to represent context exploratory behavior and escape motivation ^37–39^ and was included in addition to freezing on all days to assess active and passive behaviors at baseline and during threat evoked by CO_2_ itself or on re-exposure to context.

### Acoustic Startle

One week following the CO_2_ exposure (Fig 2 layout), startle response to an unexpected acoustic stimulus was measured using the SR-LAB startle response system (San Diego Instruments, San Diego, CA) in standard room air as previously described with modifications ^27,40^. The enclosure was of sufficient size to restrict but not restrain the animal, as it allowed mice to turn around. The chambers were calibrated using the SR-LAB standardization unit (San Diego Instruments, San Diego, CA), prior to testing. Background noise in the chamber was maintained at 68 dB. After a 5min acclimatization period, mice were exposed to 10 trials of 110 dB stimuli over the 68dB background (40 ms duration; 30-38 s inter-trial interval) followed by 30 trials included randomly generated 0, 95, 100, 110, 115 and 120db stimuli over background (40 ms duration; 30-38s inter-trial interval). Movement inside the tube was detected by a piezoelectric accelerometer below the frame. For each trial, measurements were taken at 1 ms intervals for a response window of 150 ms following the startle stimulus using National Instruments Data Acquisition Software (San Diego Instruments, San Diego, CA). The maximum response amplitude (Vmax; mV) within the recording window was used for data.

**Figure 2.**
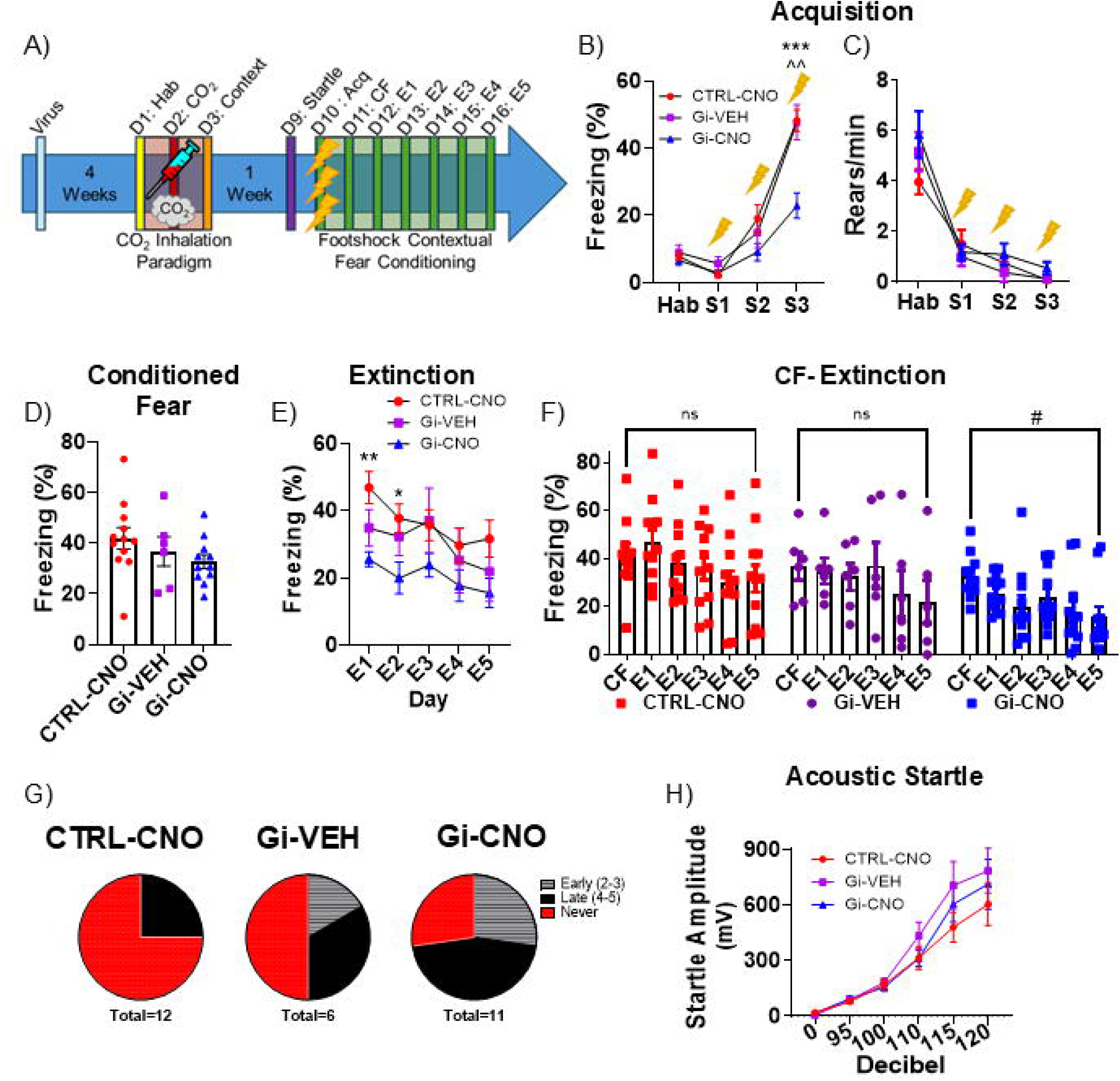
SFO-IL projection regulates delayed effects of CO_2_ inhalation on threat responding in footshock contextual fear conditioning- extinction. (A) Top panel shows the layout of the fear conditioning paradigm. Mice received a single injection of CNO or vehicle 30 min prior to CO_2_ exposure *a week* prior to testing for acoustic startle and footshock contextual fear conditioning and extinction. (**B**) CNO administration in Gi-CNO mice one week earlier significantly reduced freezing on fear acquisition day, following foot shocks versus control virus (CTRL-CNO) and Gi-vehicle (Gi-Veh) groups. (**C**) Post shock reduction in rearing behavior was similar between groups with no significant difference. (**D**) No significant difference in 24 h post-shock conditioned freezing was observed between groups (**E**) Comparison of extinction freezing over 5 days revealed significant reduction in Gi-DREADD mice compared to control virus and vehicle groups. (**F**) To assess efficacy of extinction a within group analysis of extinction freezing over days compared to conditioned freezing on day 2 (CF) revealed significantly reduced CF to extinction day 5 freezing in Gi-CNO mice, while no significant reduction in freezing was observed in vehicle and virus control groups. (**G**) Pie charts represent the distribution of mice within groups that either achieved extinction (freezing <10%) in the early phase (days 2-3), late phase (days 4-5) or never. Extinction was achieved in only 25% of CTRL-CNO mice (0% early, 25% late, 75% never) and 50% of Gi-VEH mice (16% early, 33% late, 50% never). In contrast, 72% of Gi-CNO mice achieved extinction (27% early, 45% late, 27% never). (**H**) Mean startle amplitude to a range of acoustic stimuli ranging across 95, 100, 110 and 120=:db is presented. Significant effects of decibel but no significant group differences were noted. Data are mean ± SEM. n= 6-12 mice/group * p<0.05 **p<0.01 between CTRL-CNO and Gi-CNO; ^^ p<0.01 between Gi-VEH and Gi-CNO, ^#^ p<0.05 between CF and E5; ns=not significant. Hab = Habituation; S1-S2-S3 = footshocks 1, 2, or 3; CF= Conditioned Fear; E1-E5 = extinction day 1-5.

### Contextual Fear Conditioning

Mice underwent a foot shock contextual conditioning paradigm to investigate fear acquisition, conditioned fear, extinction and reinstatement in standard room air as described in previous studies ^27,40^. Operant chambers housed in sound attenuated isolation cabinets were used (Clever Sys Inc.). The floors of the chambers consisted of stainless-steel grid bars that delivered scrambled electric shocks. The grid, floor trays and chamber walls were wiped with 10% ethanol and allowed to dry completely. Mice acclimated for 5 min before receiving 3 shocks (0.5mA, 1 sec duration, 1 min apart). Mice returned to the chamber the next 6 days and behaviors were recorded for 5 min without shocks to measure conditioned fear (Day 2) and extinction (Days 3-7). After the first 5 min on day 7, mice received 1 reminder shock (0.5mA, 1 sec) and remained in the chamber for 5 min to measure reinstatement of fear. Freezing behavior was scored using automated FreezeScan software (CleverSys Inc.). We also assessed post-shock rearing behaviors on acquisition and reinstatement days as a measure of active defensive behavior. Frequency of rearing was scored by a trained observer blinded to experimental condition.

### Water consumption following 24h Dehydration

Water was removed from mouse cages for 24 h. Mice were then placed in individual cages and received injections of CNO (3mg/kg) or vehicle. After 30 m, mice received water sippers^41^. Water consumption was assessed at 15 m, 30 m, 60 m and 120 m after which time they were returned to their home cages.

### HPA quantification following Restraint Stress

Mice were placed into individual cages and received injections of CNO (3mg/kg) or vehicle. After 30 m, mice were placed into restrainers and baseline blood samples were immediately collected by quickly clipping the distal tip of the tail and collecting ∼20ul of blood into EDTA coated microcapillary tubes. After 30 m, an additional blood sample was collected, and mice were removed from the restrainers. A final blood sample was collected 2 h after the beginning of the restraint. Blood samples were left on ice until being spun at 1000 rpm for 15 m at 4°C. Plasma was collected and frozen until corticosterone (cort) concentration was quantified by radioimmunoassay as previously described ^42^ using 125I RIA kits (MP Biomedicals Inc., Orangeburg NY).

### Optogenetic Stimulation of SFO

Animals were habituated to the attachment of the fiberoptic cable (Plexon, 94021-050) to the fiberoptic probe implant (Plexon, 94091) for 20 m while no light was administered. Following reattachment 24h later, animals were exposed to 473nm LED pulses (20ms) at 20Hz delivered through the fiberoptic cable (Plexon). The output was maintained at 19.0mW as measured at the tip of the fiber cable. Animals were stimulated for 20s (ON phase), followed by absence of light for 10 s to minimize neuronal excitotoxicity (OFF phase). This 20 s-ON, 10 s-OFF alternating pattern was repeated for 10 m, as described before for optogenetic activation of the SFO ^43^. After 90 m, animals were sacrificed by transcardial perfusion and tissue was processed for cFos immunohistochemistry (IHC) as described below.

### Fiber Photometry Recordings and Analysis

A commercially available photometry hardware and software package (Tucker Davis Technologies, Alachua, FL) was used for IL recordings during CO_2_ inhalation. To transmit light and record GCaMP6f fluorescent signal, we used a RZ10X processor and amplifier emitting LED light to excite GFP and isosbestic control (465 nm and 405 nm, respectively) delivered to two minicubes (Doric Lenses, Quebec, Canada). The signal was processed with Synapse software suite (Tucker Davis Technologies, Alachua, FL). LEDs were powered with sufficient mV to reach 30uW. Prior to recordings, fiberoptic cannula were photobleached for 4h. The fiber optic implant was coupled to the optical patch clamp (Ø 400 µm, NA 0.37, Hytrel loose tube 1.1 mm jacket, Doric Lenses, Quebec, Canada) using a 1.25 mm zirconia mating sleeve (Doric Lenses) then mice were placed in the CO2 chamber and allowed to habituate for 5 minutes. Breathing air was then infused into the chamber for 10m to establish baseline activity, followed by 5% CO_2_ for 10m. Photometry data were extracted and analyzed using a Python-based graphical user interface (GUIs): GuPPY^44^. The data was first filtered by a zero-phase moving average linear digital filter before the ΔF/F was calculated using a fitted control isobestic signal using a least squares polynomial fit of degree. The ΔF/F was used to assess the change in amplitude and frequency of transients over time using a 15-sec moving window and the transient frequency and amplitude was averaged over 2 min bins before being plotted using graphpad (GraphPad Software Inc., San Diego, CA).

### Immunohistochemistry (IHC) and Image analysis

Mice were transcardially perfused with 4% paraformaldehyde and processed for IHC as previously described ^24,27^. Briefly, 30 μm coronal brain sections were cut and stored in cryoprotectant (0.1M phosphate buffer, 30% sucrose, 1% polyvinylpyrrolidone, and 30% ethylene glycol) at −20°C. For immunolabeling, slices were washed 5 times for 5 minutes (5 x 5m) in PBS, incubated with 3% H_2_O_2_ for 10 min then washed (5 x 5m) in PBS. Tissue was then incubated in blocking solution for 1h. To track GFP immunofluorescence to label the retrogradely transported cre virus slices were incubated overnight with primary antibodies against green fluorescent protein (GFP; 1:1000 ThermoFisher #A6455). The following day, sections were washed again (5 x 5m) in PBS then were incubated in Alexa488 secondary antibody (Alexa488 anti-Rb 1:500 Jackson Immuno #711-545-152) for 1h. They were then washed again (5 x 5m) in PBS and incubated in blocking solution for 1h. To label the DIO-DREADD-Gi mCherry and DIO-mCherry (RFP) viruses, tissue was then incubated overnight with primary antibodies against red fluorescent protein (RFP; 1:1000 Rockland cat#600-401-379). The following day, sections were washed again (5 x 5m) in PBS then incubated in Cy3 secondary antibody (Cy3 anti-Rb 1:500 Jackson, cat#711-165-152) for 1h, washed (5 x 5m) in PB, mounted, and cover-slipped using Gelvatol (Sigma-Aldrich cat#10981-100ml). Immunolabeled sections were mounted and imaged at 20x using an AxioImager ZI microscope (Axiocam MRm camera and AxioVision Release 4.6 software; Zeiss). Subjects were delineated as “hits” if SFO (AP -0.10 to -0.80 mm) cells co-expressed RFP and GFP, and if the IL (AP 1.98 to 1.34 mm) or BNST (AP 0.50 to -0.02 mm) expressed GFP.

### Statistical Analysis

For behavioral experiments, n=12-18/group underwent surgeries, hits/misses were assessed, and mice were excluded if they did not co-express GFP and RFP in the SFO. For the SFO to IL study, final group numbers were n=12/18 (Ctrl-CNO), 8/12 (Gi-VEH) and 11/18 (Gi-CNO). For the SFO to BNST study, final group numbers were n=8/12 (Ctrl/CNO), 4/12 (Gi/VEH) and 7/12 (Gi/CNO). For the AT1R-expressing SFO-IL study, final group numbers were n=7/12 (Ctrl) and n=8/12 (Gi). Data are represented as means ± standard error and statistical analyses included students’ unpaired t-test, two-way ANOVA or three-way repeated measures ANOVA as appropriate. Welch’s corrections or Kruskall-Wallis tests were applied when distributions had unequal variance. Non-parametric Mann-Whitney tests were applied to distributions failing normality as determined by the Kolmogorov-Smirnov test. For *post hoc* analyses, Bonferroni, Dunnett or Dunn’s tests were used to correct for multiple comparisons. Grubbs’ tests were performed to determine and remove any outliers (maximum 1/group). Results were considered statistically significant at the *p*<0.05 level and statistical analyses were performed in *Prism*, version 9 (GraphPad Software, Inc., La Jolla, CA).

## Results

### 1. SFO to IL projection regulates defensive behaviors to interoceptive threat, CO_2_ inhalation

Our previous studies supported a role of the SFO in CO_2_-chemosensing and associated defensive behaviors ^24,34,45^ . Given the role of the IL in threat appraisal and reported SFO-to-IL projections ^26,46^ we tested whether this circuit is recruited in defensive behaviors during CO_2_ inhalation (schematic Fig 1A, C). Following a 4 week recovery from surgical manipulations, no group differences were observed in freezing (F_(2,27)_ = 0.137, p=0.872, Fig 1D) or rearing (F_(2,29)_ = 2.200, p = 0.129, Fig 1E) behaviors while mice habituated to the CO_2_ inhalation chamber. On day 2, behaviors during CO_2_ inhalation (30 min post veh/CNO injection) were assessed. A significant group difference in CO_2_-evoked freezing behavior was observed (F_(2,29)_ = 4.146, p = 0.027, Fig 1F). Post hoc analysis revealed a significant reduction in freezing in the CNO treated- inhibitory DREADD Gi receptor group (Gi-CNO) compared to Gi-DREADD mice receiving vehicle (Gi-VEH) (p<0.05). Interestingly, SFO-IL circuit inhibition resulted in increased exploratory rearing behavior (F(_2,28)_ = 3.412, p = 0.0483, Fig 1G) suggesting that SFO-IL circuit inhibition induces a shift in defensive strategy adopted during interoceptive threat, CO_2_. Post hoc analysis revealed significantly higher rearing in Gi-CNO mice compared to control mCherry-CNO treated mice (CTRL-CNO; p<0.05). In addition to evoking spontaneous responses during inhalation, CO_2_ also acts as an unconditioned stimulus to produce conditioned contextual freezing specific to exposure context^24,32^, a response relevant to avoidance of panic contexts in panic patients due to associative conditioning ^47^. Re-exposure to the CO_2_ testing context on day 2 evoked freezing in control virus and vehicle groups, an effect that was significantly attenuated in Gi-CNO mice (F_(2,27)_ = 7.284, p = 0.0032, Fig 1H). Post hoc analysis revealed that Gi-CNO mice froze significantly less (p<0.01) than both control groups. Rearing behavior during context re-exposure also revealed significant group differences (F_(2,28)_ = 5.795, p = 0.0083, Fig 1I). Consistent with increased exploration observed during day 2 CO_2_ inhalation, Gi-CNO mice exhibited significantly higher rearing upon context re-exposure (p<0.05) versus control groups, suggesting that the SFO-IL circuit also regulates context conditioned behaviors associated with the CO_2_ experience.

To confirm the specificity of the SFO-IL circuit, we investigated whether SFO projections to the BNST that have been recently reported in inflammation-induced anxiety ^25^ can regulate CO_2_-evoked defensive behaviors. Following recovery from surgical manipulations, no differences were observed in freezing (F_(2,17)_ = 0.987, p=0.393, Suppl Fig 1D) or rearing (F_(2,17)_ = 2.011, p = 0.165, Suppl Fig 1E) behaviors while mice habituated to the CO_2_ inhalation chamber. The following day, no significant group differences were observed during CO_2_ inhalation in freezing (F_(2,19)_ = 2.267, p =0.134, Suppl Fig 1F) or rearing (F_(2,19)_ = 2.477, p = 0.114, Suppl Fig 1G). Consistent with no behavioral effects of SFO-BNST circuit inhibition during CO_2_ inhalation, no significant group differences were observed when mice were re-exposed to context in the absence of CO_2_ in freezing (F_(2,17)_ = 0.006, p=0.995, Suppl Fig 1H) or rearing (F_(2,19)_ = 0.225, p=0.801 Suppl Fig 1I). Collectively, these data suggest that SFO to IL projections regulate defensive behavioral strategies adopted during exposure to interoceptive threat, CO_2_ and associative context conditioned behaviors.

### 2. SFO-IL projection regulates delayed effects of CO_2_ inhalation on contextual fear extinction

In addition to acute aversive effects during inhalation, CO_2_ impacts delayed emotional reactivity to subsequent traumatic experiences in veterans ^8^. As previously reported by us, exposure of CO_2_ mice to footshock contextual fear conditioning-extinction one week later showed enhanced contextual fear and delayed extinction learning ^27,34,36^. In addition to sensing CO_2_ induced acidosis ^24^, the SFO also detects other aversive interoceptive signals. We examined whether dehydration, an aversive homeostatic threat sensed by the SFO to regulate drinking behaviors, may impact fear conditioning and extinction a week later (Suppl Fig 2). Mice with a prior history of acute 24h dehydration (physiological stressor that promotes reduced body weight and food intake Suppl Fig 2B,C) elicited comparable freezing to the control group during fear conditioning and extinction. For fear acquisition (Suppl Fig 2D) there was a significant main effect of time (F_(1.551, 31.01)_ = 53.95, P<0.0001), but no treatment F_(1, 20)_ = 0.425, p= 0.521) or time x treatment interaction F_(1.551, 31.01)_ = 0.7891, p=0.433) effects. For contextual fear memory (Suppl Fig 2E) no significant group differences were observed (t test, t=0.2390, df=20, p=0.81). Extinction freezing (Suppl Fig 2F) was also similar between groups with only a significant effect of time (two-way ANOVA, F_(5, 121)_ = 2.838, p=0.018) but no effect of treatment (two-way ANOVA, F_(1, 121)_ = 1.614; p=0.206) or a time by treatment interaction (F_(5, 121)_ = 0.3728, p=0.866). In contrast, prior single exposure to CO_2_ a week earlier produced significant effects on fear acquisition, conditioned fear and compromised extinction. No group differences occurred in baseline freezing to the CO_2_ chamber during habituation (t_(59)_ = 0.429, p = 0.669, Suppl Fig 2G), however CO_2_ inhalation the following day significantly increased freezing (t_(59)_ = 3.129, p = 0.003, Suppl Fig 2H). Re-exposure to the CO_2_ inhalation context also significantly increased freezing (t_(57)_ = 2.734, p = 0.009, Suppl Fig 2I). During fear conditioning acquisition one week later (Suppl Fig 2J), we found significant effects of inhalation (F_(1,60)_ = 6.518, p = 0.0132), time (F_(2.227, 133.6)_ = 160.3, p < 0.0001), and time x inhalation (F_(2.227, 133.6)_ = 3.033, p = 0.046). Post hoc tests revealed prior CO_2_ inhalation significantly increased freezing to the 2^nd^ shock (p<0.05). When returned to the fear conditioning context the following day (Suppl Fig 2K), mice who underwent CO_2_ inhalation one week prior to fear acquisition froze significantly more than air exposed mice (t_(58)_ = 3.456, p = 0.001). During extinction (Suppl Fig 2L), freezing to the context was also significantly higher in CO_2_-exposed mice (inhalation: F_(1, 58)_ = 9.003, p = 0.004; time: F_(2.814, 163.2)_ = 30.56, p < 0.0001 but no interaction F(4,232) = 1.056, p = 0.379). Posthoc tests revealed significantly higher freezing in mice with prior CO_2_-exposure on days 1, 3 and 5 (p < 0.05). Thus, fear memory modulation by aversive interoceptive threats does not appear to be a generalized phenomenon.

Given altered threat responding during CO_2_ inhalation in mice with SFO-IL circuit inhibition we next examined whether not having the circuit online during CO_2_ could impact delayed CO_2_ effects on fear conditioning-extinction. Intriguingly, even though SFO-IL circuit inhibition was undertaken 7d earlier, Gi-CNO mice elicited reduced freezing on exposure to footshocks during fear acquisition (Fig 2B). A two-way ANOVA revealed a significant overall effect of treatment (F_(2,27)_ = 9.154, p = 0.001), time (F_(3,27)_ = 102.5, p < 0.0001) and time x treatment interaction (F(_6, 81)_ = 6.050; p < 0.0001). Post hoc analysis revealed significantly reduced freezing following the 3^rd^ shock in the Gi-CNO group compared with both control groups (p<0.05). Interestingly, all groups elicited comparable reduction in rearing behavior (F_(2,27)_ = 9.638, p=0.001, Fig 2C) that is observed in threatful contexts. To assess whether reduced post shock freezing reflects compromised fear learning in Gi-CNO mice we assessed contextual freezing 24 h later Gi-CNO mice elicited comparable freezing on re-exposure to the shock context (F(_2,26_) = 1.522, p=0.237, Fig 2D) indicative of normal associative fear memory in these mice, despite a distinct defensive behavioral response during footshock exposure. We further examined the long-term effects of prior SFO-IL circuit manipulation during CO_2_ on contextual fear extinction, Gi-CNO mice froze less than control groups during extinction (Fig 2E). There was a significant overall effect of treatment (F_(2,26)_ = 3.697, p=0.039) and time (F_(4,26)_ = 8.878, p<0.0001), but no time x treatment effect (F_(8,104)_ = 0.929, p=0.496). Post hoc tests revealed significantly reduced freezing in Gi-CNO mice compared to the (CTRL-CNO) group on days 1 (p<0.01) and 2 (p<0.05). We further compared the efficiency of extinction by comparing freezing on conditioned freezing on day 2 (CF) with freezing throughout extinction within each group (Fig 2F). Within group analysis showed a significant reduction in freezing between CF and extinction day 5 (E5) only in the Gi-CNO group (p<0.05). No significant difference in freezing from CF through extinction was observed for the control groups indicative of a CO_2_-associated extinction deficit consistent with our previous studies ^27,34^. We further assessed distribution of mice within each group that achieved extinction (freezing <10%) in the early phase (days 2-3), late phase (days 4-5) or never (Fig 2G). We found 72% of Gi-CNO mice achieved extinction (27% early, 45% late, 27% never). In contrast, extinction was achieved in only 25% of CTRL-CNO mice (0% early, 25% late, 75% never) and 50% of Gi-VEH mice (16% early, 33% late, 50% never).

To investigate specificity of SFO-IL circuit in regulating delayed effects of CO_2_ on fear conditioning-extinction we hypothesized that modulation of SFO-BNST projections (that did not significantly impact behavior during CO_2_ inhalation or context re-exposure) would not regulate delayed CO_2_ effects a week later. Consistent with this, no significant group differences were noted in all phases of fear conditioning in SFO-BNST CO_2_ exposed mice. During fear acquisition (Suppl. Fig 1J), there was an overall effect of time (F_(3,51)_ =14.78, p<0.001), but no effect of treatment (F_(2,17)_ =0.078, p=0.925) or time by treatment interaction (F_(6, 51)_ =0.107, p=0.995). Context conditioned freezing 24h later (Suppl Fig 1K) also showed no treatment effect (F_(2,16)_ =0.638, p=0.542). For extinction (Suppl Fig 1L), there was a significant overall effect of time (F_(4,64)_ =7.096, p=0.003), but no significant treatment (F_(2,16)_ =0 .143, p=0.868) nor time by treatment interaction (F_(8, 64)_ = 0.606, p=0.770). Collectively, these data suggest that in addition to regulating defensive behaviors during CO_2_ inhalation, the SFO-IL projection impacts delayed effects of CO_2_ on defensive responding to distinct exteroceptive footshock stimulus and extinction of fear.

### 3. Threat regulation by the SFO to IL projection does not generalize to other aversive experiences

We questioned whether the SFO-IL circuit would also regulate response to other threat exposures of distinct modalities: sensory (acoustic), homeostatic (thirst) and psychogenic (restraint). On exposure to an acoustic startle stimulus of increasing decibel intensity all mice significantly increased their startle response (F_(5, 135)_ = 70.52; p<0.0001, Fig 2H), however, no significant group differences were observed in the startle response amplitude (F_(2,27)_ = 0.8685; p=0..869) or a treatment by decibel interaction (F_(10,135)_ = 0.869; p=0.564).

The SFO has previously been implicated in psychogenic stress-associated hypothalamic pituitary adrenal (HPA) response ^48,49^ and the IL regulates acute stress-induced neuroendocrine response^50^. To determine whether SFO-IL projections regulate acute psychogenic stress induced HPA response, blood corticosterone levels were quantified at baseline (0m), immediately (30 min) and 120 min post restraint stress. Although there was a significant overall effect of time (F_(2,30)_ = 76.49; p<0.001, Fig 3A), no effect of treatment (F(_2,15)_ = 0.265; p=0.770) or time by treatment interaction (F_(4,30)_ = 0.781; p=0.546) was observed.

**Figure 3.**
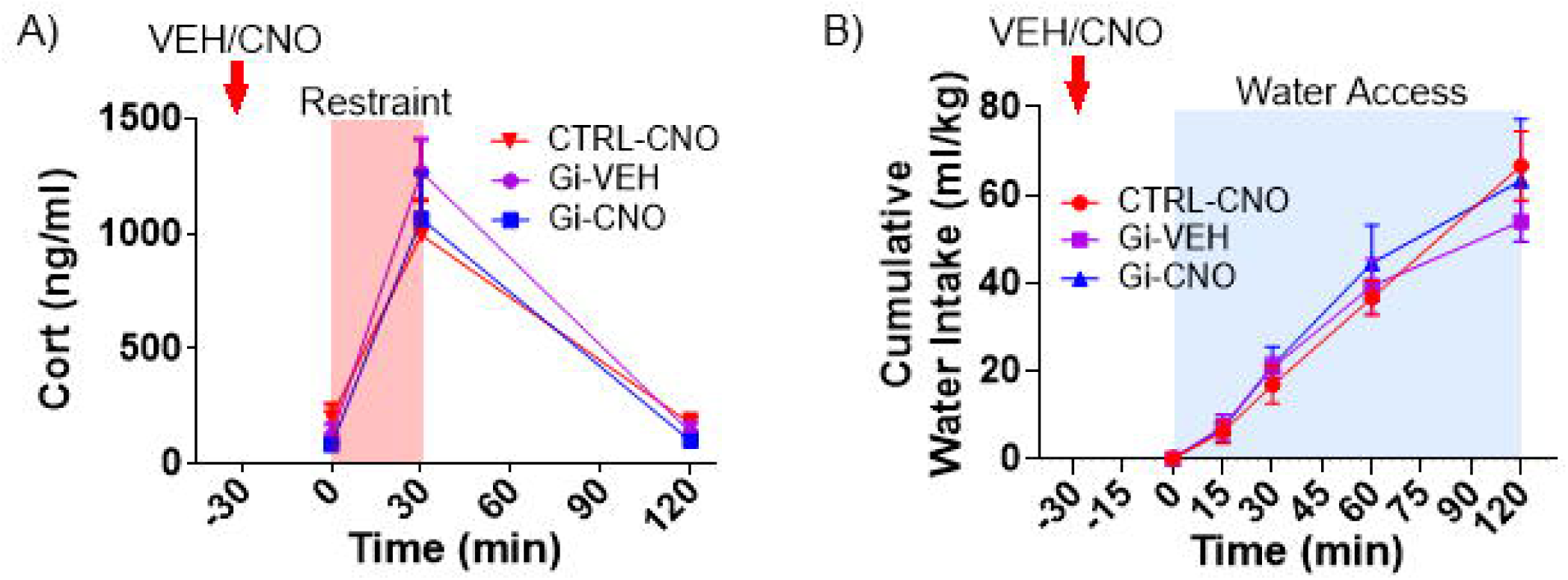
Threat regulation by the SFO to IL projection does not generalize to aversive psychogenic or homeostatic stress: (A) Acute psychogenic restraint stress induced HPA response assessed as blood corticosterone levels at baseline (0m), immediately (30 min) and 120 min post stress not impacted by SFO-IL circuit modulation. No significant difference was observed between Gi-CNO, Gi-vehicle and CTRL-CNO groups. (B) Cumulative water consumption after 24h dehydration stress was comparable between Gi-CNO and control groups as no significant group differences were observed. Represented data are mean ± sem (n= 4-12 mice/group).

The role of the SFO in sensing homeostatic fluctuations in osmolality to induce water intake is well established ^22,51^. Thus, we examined whether SFO to IL circuit regulates water consumption in a dehydrated state. Quantification of cumulative water consumption over 2h revealed a significant overall effect of time (F(_3,57)_ = 62.14, p<0.0001 Fig 3B), but no effect of treatment (F_(2,19)_ = 0.102, p=0.903) or a time x treatment interaction (F_(6,57)_ = 0.690, p=0.659). Collectively, our data suggests that the SFO-IL projection recruitment does not generalize to other threat modalities representing aversive sensory, homeostatic and psychogenic triggers.

We also assessed the effect of inhibiting SFO to BNST projections on acoustic startle, restraint stress induced HPA response and water consumption following dehydration. On exposure to an acoustic startle stimulus, all mice significantly increased their startle response with increasing decibel intensity (F_(5, 80)_ = 15.41; p<0.0001, Suppl Fig 1M), however, no significant group differences were observed in the startle response amplitude (F _(2,16)_ = 0.255; p=0.778) or a treatment by decibel interaction (F_(10,80)_ = 0.378; p=0.813). In the HPA restraint stress test, while there was a significant overall effect of time on corticosterone (Suppl Fig 1N; F_(2,20)_ =18.92, p<0.0001), there were no effects of treatment (F _(2,20)_ =0.123, p=0.885) nor time x treatment interaction (F_(4,40)_ =0.493, p=0.741). Similarly, we found an overall effect of time on water consumption (Suppl Fig 1O; F _(2,51)_ =41.88, p<0.0001), but no effect of treatment (F_(2,17)_ =0.466, p=0.636) nor time x treatment interaction (F(6,51)=1.161, p=0.342). Overall, these data suggest that SFO projections to the IL (or BNST) do not regulate acoustic startle response, post-stress neuroendocrine response or homeostatic dehydration stress induced water intake.

### 4. SFO angiotensin receptor, AT1R^+ve^ neuronal projections to the IL regulate CO_2_- associated spontaneous and conditioned fear as well as delayed effects on fear extinction

We next sought to determine what neuronal cell type within the SFO may drive CO_2_-associated defensive behaviors and delayed effects on fear memory and extinction. We hypothesized involvement of the renin angiotensin system (RAS) /angiotensin receptor type 1 (AT1R) expressing neurons. RAS polymorphisms have been reported in panic disorder^28^, and PTSD patients^29,30^ . Importantly, CO_2_ upregulates SFO AT1R expression and AT1R antagonism attenuates CO_2_ evoked freezing and associated contextual fear ^31^. Using a high throughput droplet-based-single-cell RNA sequencing dataset reported previously^52^, we mapped the expression of the AT1R transcript (Agtr1a) on SFO cell type clusters. An unsupervised clustering on graph-based representation of cellular gene expression profiles visualized in a uniform manifold approximation and projection (UMAP) embedding revealed diverse cell types characterized by unique transcriptional signatures (t-SNE plot (Fig 4A). A feature plot of cell-type specific expression of *Agtr1a* (Fig 4B) and violin plot analysis (Fig 4C) revealed a neuron-skewed expression of *Agtr1a* localized specifically to excitatory neurons. Marked co-localization of AT1R-tdtomato expression was observed with calmodulin kinase II (CAMKII ^+ve^ neurons (Fig 4D) that are primary SFO projection neurons^53^. Collectively, these observations support that AT1R^+ve^ SFO neurons represent excitatory projections. Using a cell-type specific intersectional chemogenetic approach, we examined whether inhibition of AT1R^+ve^ -SFO to IL projection regulates CO_2_-associated defensive behaviors (Fig 4E). AT1R-cre mice received a viral infusion of a cre-dependent flpase into the SFO and bilateral injections of a flp-dependent, retrogradely transported inhibitory DREADD into IL (Fig 4F,G). After recovery they underwent the CO_2_-fear conditioning paradigm (Fig 4E). No significant behavioral effects on freezing (t(13) = 0.106, p = 0.917; Fig 4H) and rearing (t(13) = 0.077, p = 0.940; Fig 4I) were observed during habituation Consistent with our hypothesis, CNO administration in Gi-DREADD AT1R-Cre mice 30 min prior to CO_2_ inhalation showed a significant reduction in freezing (t(13) = 2.937, p=0.012, Fig 4J) and increased rearing behavior (t(13) = 3.990, p=0.002; Fig 4K;) during CO_2_ inhalation. Re- exposure to the inhalation context 24h post CO_2_ showed no significant group differences in freezing (t(12)=1.146, p=0.274 Fig 4L) or rearing (t(13)=0.593, p=0.564 Fig 4M), likely due to poor CO_2_ context conditioned behaviors in the background strain (C57BL6) of AT1R-Cre mice ^36^. Interestingly, exposure of control and Gi-DREADD mice to a footshock contextual fear conditioning paradigm revealed long term behavioral effects of circuit manipulation during prior CO_2_ inhalation. Exposure to footshocks during fear acquisition revealed an overall effect of time (F_(3,39)_ =36.38, p<0.001, Fig 4O), but no effect of treatment (F(_1,13)_=0.451, p=0.514) or a time x treatment interaction (F _(3,39)_ =0.545, p=0.545). Re-exposure to the shock context revealed no significant effect of treatment on contextual conditioned fear ( t(12)=0.929, p=0.371, Fig 4P). Exposure to the context for the next 5 days to assess extinction of fear revealed significantly reduced extinction freezing in mice with inhibition of AT1R^+ve^ SFO-IL projection during CO_2_ inhalation a week earlier (Fig 4Q). A two-way ANOVA showed a significant overall effect of treatment (F_(1, 12)_ =5.641, p=0.035) and time (F _(4,48)_ =6.735, p<0.001), but no significant time x treatment interaction (F_(4,48)_=0.963, p=0.437). Post hoc analysis revealed a significant reduction in extinction day 4 freezing in Gi-DREADD mice. Comparison of extinction freezing across days with conditioned fear freezing (Fig 4R) within groups revealed minimal extinction learning in CO_2_ exposed control mice with normal AT1R SFO-IL circuit signaling as no significant differences were observed compared to conditioned fear freezing. In contrast CO_2_-Gi-DREADD mice elicited significantly lower freezing on extinction days 3 (p<0.001) day 4 (p<0.001) and day 5 (p<0.01) indicating improved extinction in these mice. Thus, CO_2_ associated extinction deficits require AT-1R^+ve^ neuronal projection from the SFO to the IL cortex.

**Figure 4.**
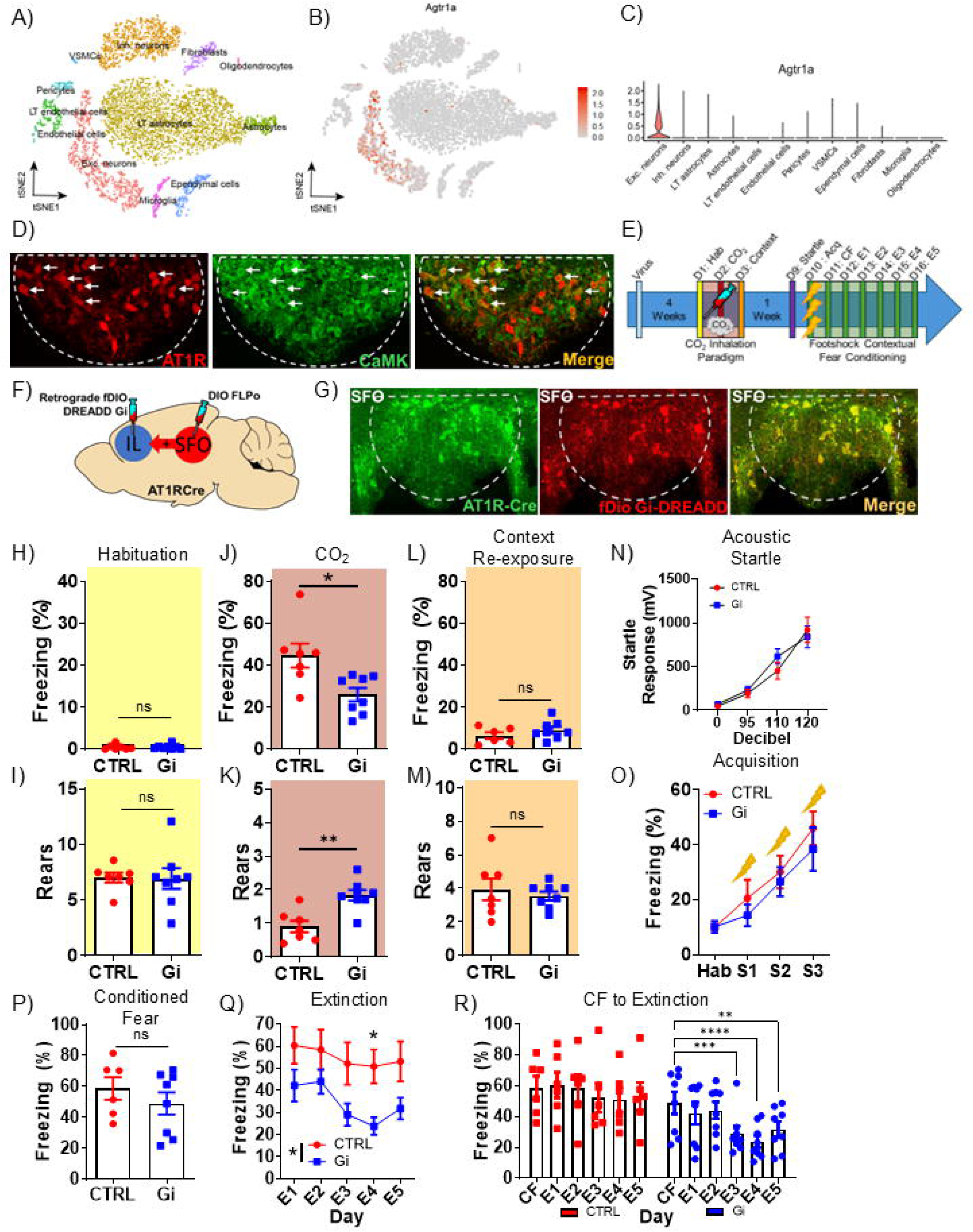
SFO angiotensin receptor, AT1R^+ve^ neuronal projections to the IL regulate CO_2_- associated spontaneous and conditioned fear as well as delayed effects on fear extinction. **(A)** Single-cell RNA-sequencing analysis showing cellular diversity in the SFO with 13 transcriptomic cell classes. Data shown in a tSNE embedding of 7,950 cells with color-coded cell identity. **(B)** Log-normalized expression of angiotensin receptor type 1A transcript (Agtr1a) in SFO cell classes. **(C)** Violin plot of log-normalized expression of Agtr1a in SFO cell classes highlighting expression in excitatory neurons. Data in A-C reanalyzed and plotted from a previous study (Pool et al, 2020) [50]. (**D**) SFO AT1R^+ve^ neurons (AT1aR-tdTomato reporter mice) show co-localization with calcium/calmodulin-dependent protein kinase (CaMK). (**E, F**) Intersectional chemogenetic strategy and experimental layout. A single injection of CNO was administered 30 min prior to CO_2_ inhalation (see text for details). (**G**) ) Images from the SFO show Cre-expressing AT1R neurons (green) and FlPo-dependent Gi-DREADD mcherry (RFP) expressing SFO to IL neurons (red) and colocalized cells in merged image. For habituation, no significant group difference was observed in freezing (**H**) or rearing (**I**). During 5% CO_2_ inhalation Gi-CNO mice elicited a reduction in freezing (**J**) and increase in rearing (**K**) compared to control (CTRL) group. On context re-exposure for conditioned behaviors no significant group difference was observed in freezing (**L**) or rearing (**M**). Mean startle amplitude to a range of acoustic stimuli ranging across 95, 100, 110 and 120[db is presented (**N**) Significant effects of decibel but no significant group differences were noted. (**O**) Freezing on fear acquisition day was similar between Gi-CNO and control (CTRL) groups. (**P**) No significant difference in 24 h post-shock conditioned freezing was observed between groups (**Q**) Comparison of extinction freezing over 5 days revealed significant reduction in Gi-DREADD mice compared to control group. To assess efficacy of extinction a within group analysis of extinction freezing over days compared to conditioned freezing on day 2 (CF) (**R**) revealed significantly reduced CF to extinction freezing in Gi-CNO mice, while no significant difference in freezing was observed in the control group. Data are expressed as mean ± sem (n= 7-8 mice/group) * p<0.05 **p<0.01 ***p<0.001 ****p<0.0001 ns=not significant.

To assess whether delayed effects on extinction in Gi-DREADD mice would generalize to other aversive exposures, we exposed Gi-DREADD mice with CNO administered prior to CO_2_ inhalation to varying acoustic startle stimulus. While a significant overall effect of decibel intensity was observed (F_(3,36)_ = 48.26, p<0.001, Fig 4N), there was no significant effect of treatment (F_(1,12)_ = 0.209, p=0.656) or a treatment x decibel interaction (F_(3,36)_ = 0.898, p=0.452).

### CO_2_-inhalation and SFO modulate infralimbic neuronal activation

Currently, there is no information on the effects of CO_2_ inhalation or the SFO on IL neuronal activation. Exposure of FosCreERT2.Ai9 mice to CO_2_ inhalation (Fig 5A) led to significant reduction in tdTomato^+ve^ cells in the IL cortex (t(6) = 2.986, p = 0.03, Fig 5B,C). Importantly, tdTomato^+ve^ cell counts showed an inverse correlation with freezing (r^2^ = - 0.69, p=0.02, Fig 5D) and positive correlation with rearing (r^2^ = - 0.47, p=0.08, Fig 5E) suggesting a potential involvement of the IL in defensive behaviors during inhalation. Consistent with our behavioral findings no significant difference in tdTomato^+ve^ cell counts was observed in the BNST (t(6) = 0.440, p=0.67, Suppl Fig 3B) and there was no correlation of trapped cells with freezing (r^2^ = - 0.051, p=0.62 Suppl Fig 3C) or rearing (r^2^ = - 0.07, p=0.56 Suppl Fig 3D). To determine neuronal response during CO_2_ inhalation we recorded pre and post CO_2_ changes with IL-targeted infusion of a CaMKII promoter driven GCaMP6f and IL-implanted fiberoptic cannula (Suppl Fig 3E). Analysis for baseline vs CO_2_ revealed a significant reduction in mean transient amplitude (One way ANOVA F(5, 257) = 14.82, p<0.001; Suppl Fig 3F) and a trend for a reduction in frequency (One way ANOVA F(5,257) = 2.164, p= 0.066; Suppl Fig 3G) consistent with reduced CO_2_-evoked IL excitation. To determine whether activation of the SFO could impact IL neuronal activation, we delivered channelrhodopsin (ChR2) targeted to SFO-CAMKIIa expressing neurons (Fig 5F). Optogenetic activation of SFO excitatory projection neurons resulted in reduced cFos in the IL, particularly in the deep layers 4-6 in ChR2 mice (t test, total t(8) = 1.934; p=0.08, layers 4-6 t(8) = 2.614, p= 0.03, layer 1-3 t(8) = 0.581, p = 0.57; Fig 5G). The role of IL inhibitory microcircuits in regulation of fear expression is well established^61,62^. Synchronization of firing of mPFC projection neurons is regulated by GABAergic interneurons, ^54,55^ and parvalbumin (PV) and somatostatin (SST) subtypes have been implicated in fear regulation ^54^. Quantification of Fos /PV^+ve^ and Fos /SST^+ve^ cells in ChR2/eGFP mice revealed a significant increase in Fos/PV^+ve^ cells in IL layers 4-6 (t(8) = 4.538, p= 0.001; Fig 5H,I) but not the PL (t(8) = 0.55, p= 0.59; Fig 5J), while no differences were noted in Fos /SST^+ve^ cells in IL (t(7) = 0.61, p= 0.55; or PL (t(7) = 0.16, p= 0.87; Suppl Fig 3H-J). Lastly, we examined IL PV interneurons post CO_2_-FC behavior (Fig 5K). A significant increase in IL Fos/PV^+ve^ cells were observed in CO_2_ exposed mice (t(14) = 2.383, p= 0.03, Fig 5L). Interestingly, Il Fos /PV^+ve^ cells showed a significant positive co-relation with freezing behavior (r^2^ = 0.36, p=0.01; Fig 5M) and inverse correlation with rearing behavior (r^2^ = 0.37, p=0.008; Fig 5N) observed during inhalation. Collectively, these data indicate that CO_2_ inhalation and the SFO can compromise IL excitation via increased activation of parvalbumin inhibitory interneurons.

**Figure 5.**
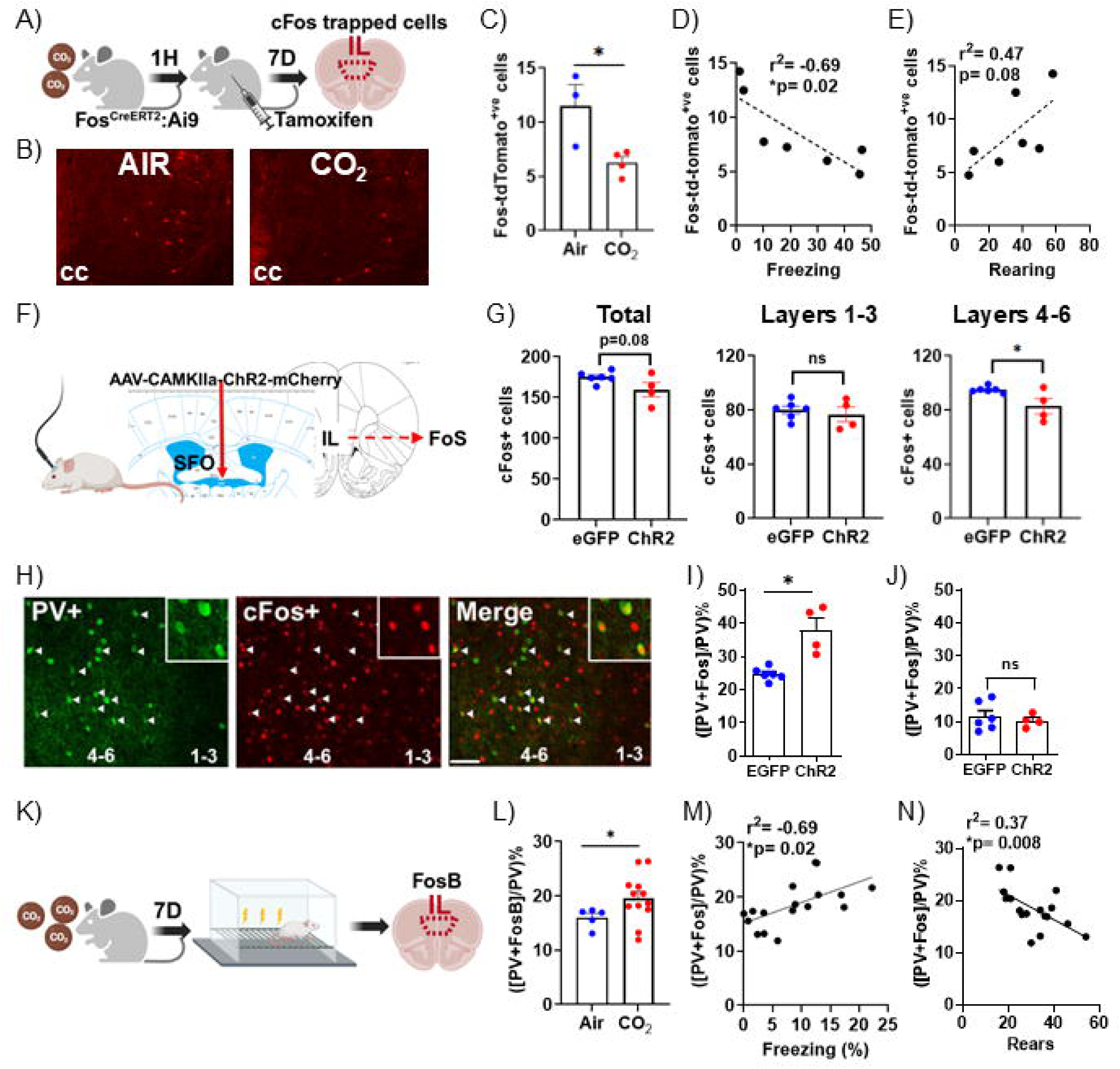
CO_2_-inhalation and SFO modulate infralimbic neuronal activation. **(A)** Layout for experiment on CO_2_ or air inhalation in Fos ^CreERT2.Ai9^ mice. Tamoxifen was injected 1 hour after inhalation. (**B**) Image panels show td-tomato^+ve^ cells in the infralimbic cortex, (**C**) Bar graph shows significantly reduced of Fos/td-tomato^+ve^ cells in CO_2_ exposed mice. Fos/td-tomato^+ve^ cells showed an inverse correlation with freezing (**D**) and positive correlation with rearing (**E)** behaviors. (**F**) Optogenetic activation of SFO CAMKII expressing neurons results in reduced cFos in IL layers 4-6 in ChR2 mice versus controls. Increased cFos + parvalbumin (PV) co-labeled neurons in the IL of ChR2 mice vs control mice (**G,H**) but not in the PL (**I**). Schematic shows layout of post behavior assessment of ΔFosB (FosB) and PV co-labelled neurons in the IL. Mice were exposed to air/CO2 inhalation followed by fear conditioning-extinction 7 days later. Tissue was collected 24h post behavior to assess ΔFosB, a marker of persistent neuronal activation. (**K**) Significantly higher PV+ FosB co-labelled neurons were observed post-behavior in CO_2_ mice versus the air group. PV+ FosB cell counts showed a positive co-relation with freezing (L) and negative co-relation with rearing (M) during inhalation. Data are expressed as mean ± sem (Fig5A-E n=3-4 mice/group; Fig5F-J n=4-6 mice/group; Fig5K-N n=5-12 mice/group) (* p<0.05; ns=not significant ).

## Discussion

Behaviors are shaped by the internal homeostatic milieu of an organism often referred to as the interoceptive state and recent work highlights interoceptive mechanisms in emotional processing and mental health ^56,57^. Currently, afferent cell-circuit pathways by which interoceptive signals modulate emotional processing are not well understood. Here, we report that CO_2_ inhalation, an aversive, interoceptive threat, engages the infralimbic cortex via afferents from the SFO, a circumventricular region critical for monitoring the internal milieu. SFO-to-IL (but not SFO-to-BNST) projections regulated defensive behaviors during CO_2_ inhalation and associative contextual fear. Notably, the SFO-IL circuit modulated CO_2_ effects on fear conditioning-extinction, but not startle, neuroendocrine response or motivated behaviors. We also established more specifically that SFO angiotensin II receptor type-1 (AT-1R)^+ve^ neuronal afferents to the IL regulate CO_2_-associated fear and the associated lasting deficits in contextual fear extinction.

The infralimbic ventromedial subdivision of the prefrontal cortex plays an important role in threat appraisal and emotional regulation ^58^, however, its role in processing aversive interoceptive threat signals is not well understood. As a visceromotor area of the cortex the IL regulates physiological responses such as gastric motility ^59^, heart rate^60^, and blood pressure^61^ via viscero-sensory nodes such as the nucleus of solitary tract and vaso-vagal connections ^62^. Afferent interoceptive signaling to the IL is currently understood to occur via midline thalamo-limbic nuclei ^63^, however, direct connections with interoceptive nodes has not been established. Our data reveal that aversive interoceptive signals can also be directly conveyed to the IL via the SFO, a key viscero-humoral circumventricular organ enabling body-brain communication.

CVOs are important sensory nodes of the interoceptive nervous system INS ^64^, an integrated collection of peripheral and central pathways and cortical regions^65^ The INS continuously senses changes to the internal homeostatic balance to promote adaptive behaviors. The sensitivity of interoceptive structures is characterized by blood brain barrier gaps for brain-body interface, and prominent non-synaptic signaling mechanisms^64^. Among the periphery and brain, the vagus nerve, lamina 1 spinothalamocortical afferents and brain stem sites such as the nucleus of stria terminalis and area postrema have previously been described as primary components of the INS for the genesis of feelings and emotional responses^64,66^. We now highlight the SFO as a key forebrain chemosensory site for threat sensing and cortical modulation of fear. Previously we reported SFO microglial acid sensing and neuroimmune signaling mechanisms in CO_2_ evoked spontaneous and conditioned fear associated behaviors^24,34,45^, although a downstream effector site was not established. Our current data show that the SFO engages the infralimbic cortex to regulate these behaviors.

CO_2_ inhalation was selected as the interoceptive stimulus for this study given its relevance as a pathologic marker in psychiatric conditions CO_2_, a threat to survival, evokes intense fear and can induce panic attacks in individuals with PD and PTSD ^3,5,6,67^. Inhalation of CO_2_ has been used as a biological challenge in the laboratory as it reliably induces unpleasant interoceptive reactions and self-reported symptoms that closely match panic attacks ^68,69^ . Emotional responsivity to CO_2_ inhalation is associated with the later development of PTSD symptoms in veterans^8^ suggesting a potential intersection of interoceptive signaling with subsequent stress and fear memory regulation. CO_2_ challenge is a cross-species translational tool^7^ that has been used in prediction of subjective distress, posttraumatic symptoms and fear extinction deficits in humans and rodents ^8,27,36,70–72^. Rising CO_2_ concentrations lead to acidosis, creating a state of homeostatic imbalance resulting in defensive behavioral responses ^73^.

CO_2_ freezing was attenuated, and rearing was increased in SFO-IL Gi-DREADD mice compared to control mice during CO_2_ inhalation and context re-exposure in the absence of CO_2_. The medial prefrontal cortex regulates adaptive decision making and risk assessment under challenging exteroceptive situations ^74^. Areas of the mPFC such as the IL regulate selection among different behavioral strategies including mobilization versus immobilization during imminent survival threats. The IL is reported to mediate competition between excitation versus inhibition of body movements, promoting behavioral strategies that involve mobilizing the body and suppressing strategies that involve immobilizing upon threat encounter ^75^. CO_2_ via the SFO may suppress IL-mediated mobilization (promote increased freezing) and circuit inhibition enables behaviors favoring movement (exploratory rearing). Interestingly, behavioral effects in Gi-DREADD mice were more robust during context re-exposure in the absence of CO_2_ than during inhalation, suggesting a stronger IL-regulatory influence on conditioned behaviors. Additionally, other CO_2_ sensing mechanisms such as the acid-sensing ion channel 1 (ASIC) in the amygdala ^35^, or hypothalamic CO_2_-sensitive orexin neurons^76^ may be engaged during inhalation.

Given the regulatory role of the SFO and IL in stress response and physiological homeostasis ^49,50,77^ we tested the engagement of the SFO-IL projection in distinct threat modalities. No significant group differences in restraint stress-evoked HPA response, acoustic startle, dehydration-indued water intake suggest that this projection may not generalize across threat modalities.

An intriguing finding of our study was the impact of an acute SFO-IL circuit manipulation during CO_2_ inhalation on fear conditioning extinction a week later. Previously, we^27,34^ and others^70,78^ reported an association of behaviors during CO_2_ inhalation with subsequent fear conditioning and extinction, highlighting responsivity to CO_2_ as a biomarker for fear pathology and extinction-based therapeutic approaches^71^ . Gi-DREADD mice exhibited active coping (less freezing, increased rearing) during selective phases of fear conditioning-extinction (similar to behaviors observed during prior CO_2_ and context exposure). Interestingly, this long-term modulation of threat reactivity expanded across threat modalities (CO_2_, footshock) and contexts (CO_2_ chamber, extinction context) suggesting overlapping interoceptive-exteroceptive mechanisms. Interestingly, the ventromedial prefrontal cortex is a key convergence node for regulating influence of the somatic state on shifts in approach-avoidance and emotion^79^. The SFO-IL circuit offers a novel convergence mechanism by which interoceptive states impact emotional responses. Interestingly, SFO-IL regulation of fear did not generalize across aversive somatic states as a prior experience of dehydration (that is sensed by the SFO for motivated thirst behavior^77^) did not have delayed effects on fear conditioning-extinction behavior. Additionally, defensive behaviors were only impacted during acquisition and extinction not during conditioned fear expression, again suggesting IL engagement during specific threat encounters consistent with previous observations^80^. No group differences in footshock contexual conditioned freezing indicates that contextual fear memory was not impacted by prior SFO-IL manipulation, and is consistent with a dispensable role of IL in associative fear memory in footshock fear conditioning ^81^. Importantly, CO_2_-Gi-DREADD mice showed improved extinction compared to CO_2_-vehicle and CO_2_-virus controls that elicited compromised extinction consistent with delayed CO_2_ effects on extinction reported by us ^27,34^ and others ^70,78^.

Our study indicates that SFO-IL mediated regulation of CO_2_-associated fear behavior may be selective. In contrast to the SFO-IL projections, CO_2_ does not appear to engage the SFO-BNST circuit to regulate spontaneous or delayed CO_2_ effects on fear. The BNST regulates anxiety and fear ^82^ to exteroceptive threats. Additionally, SFO-BNST projections also respond to interoceptive signals such as hypernatremia ^77^ and regulate inflammation associated anxiety ^25^ . Collectively, these observations indicate that SFO projections to discrete forebrain sites may regulate adaptive behaviors to distinct interoceptive signals underscoring the SFO as a multimodal sensory locus for body-brain communication.

We further identified the phenotype of SFO neurons that project to the infralimbic cortex for regulating CO_2-_ associated defensive fear behaviors. Inhibition of SFO angiotensin II receptor type -1 (AT-1R)^+ve^ neuronal projections to the IL attenuated freezing and increased rearing behavior during CO_2_ inhalation as well as improved extinction following fear conditioning a week later. Previous studies reported CO_2_-induced upregulation of SFO AT1 receptors and importantly, attenuated CO_2_-induced freezing by AT1R antagonist losartan^31^. Associations between the renin angiotensin system and fear-associated disorders have been previously reported ^29,30,83^. Retrospective longitudinal studies report improved PTSD symptoms in individuals taking ACE/angiotensin receptor blockers, like losartan^44,45^, although clinical trials have not shown efficacy ^84^. Previous preclinical and clinical studies report AT1Rs in the encoding of aversive stimuli and threat reactivity using unpleasant exteroceptive triggers ^85–87^. Although AT1Rs are abundantly expressed in multiple brain areas involved in threat, stress and fear regulation (ref), cellular mechanisms and associated circuits remain elusive. The SFO is enriched in renin angiotensin system (RAS) mediators^20^ and previous work highlighted SFO-AT1R-mediated regulation of autonomic response and motivated behaviors such as thirst and salt intake ^22,88^. Divergent downstream AT-1R^+ve^ SFO projections to the organum vasculosum of the lamina terminalis (OVLT) or the BNST regulate thirst or salt appetite, respectively ^22^. We now report that a subpopulation of AT-1R^+ve^ SFO neurons project to the IL cortex and regulate defensive behaviors to fear evoking threats such as CO_2_ but are not engaged in motivated behaviors such as thirst.

Observed CO_2_-associated fear behaviors possibly arise due to CO_2_ induced / SFO mediated dampening of the IL, a conclusion supported by our data showing reduced IL neuronal activation following CO_2_ inhalation as well as optogenetic activation of SFO neurons. Individuals with a dysfunctional vmPFC experience emotional rigidity or the inability to switch or regulate emotional responses and Impaired functioning of the ventromedial cortex and extinction deficits are consistently observed across threat and anxiety associated disorders ^2,89–91^. Hypoactivity within the IL is observed in threat-associated disorders, particularly PTSD ^90^. Consistent with our behavioral data, CO_2_ exposure and optogenetic activation of the SFO reduced IL neuronal activation. Shifts in IL activation are observed following aversive exteroceptive exposures such as chronic stress^92^ and traumatic injury^93^ that attenuate IL activity leading to cognitive impairments and fear^92,93^. Interestingly, correlation of IL FoS-tdTomato^+ve^ cells in CO_2_ exposed TRAP mice with CO_2_ freezing and rearing supports their recruitment in these defensive behaviors, an association absent in the BNST. SFO-ChR2 activation increased the proportion of PV^+^ cells in the IL but not PL, and increased PV^+ve^ / FoS^+ve^ cell counts in CO2 mice post extinction positively co-related with CO_2_ freezing and negatively with rearing. The role of IL inhibitory microcircuits in regulation of fear expression is well established ^54,55^ . Consistent with our findings footshock stress engages PV^+ve^ IL neurons and their excitation impairs delayed extinction ^94^ . Our data suggest IL PV-interneurons are receptive to afferents arising from chemosensory nodes and may serve as a cellular substrate for interoceptive regulation of fear, although exact cellular mechanisms need to be established.

Although our data highlights an interoceptive afferent cortical circuit for fear regulation, there are limitations. Our CO_2_ response readouts were restricted to behavioral endpoints. In addition to intense fear, CO_2_ induces increased autonomic and ventilatory responses ^95^. Given regulatory effects of the SFO and IL in physiological regulation ^96,97^ it would be important to assess these endpoints. Our study provides strong foundational evidence supporting SFO engagement with IL PV neurons, however, future studies are required to establish cellular mechanisms. Lastly, for consistency with our previous findings ^34,45,98^ the current study used male subjects. Recently we reported divergent, sex-dependent CO_2_ behaviors and engagement of brain stem mechanisms in female mice^36^. Given evidence of interoceptive sensitivity in women, it would be important to conduct cell-circuit studies in female subjects.

## Conclusions and Translational relevance

Ventromedial cortical deficits are associated with several psychiatric conditions. Our data reveal that interoceptive threat sensing nodes have access to the infralimbic cortex (ventromedial area 25 in humans) for behavioral regulation. Such interoceptive pathways may exist to dampen mobilization in the face of imminent danger – threat to survival as that signaled by CO_2_ to channel resources towards physiological homeostasis. Furthermore, we highlight the association of behaviors during interoceptive experiences with cortical performance in other threat scenarios; emphasizing the utility of interoceptive CO_2_ sensitivity as a biomarker for predicting psychiatric risk, as well as non-response in cortical targeted therapies^71^. Lastly, easy accessibility of BBB-compromised CVOs such as the SFO opens novel avenues for cortical therapeutic targeting. Collectively, we report a unique “bottom-up” cell-circuit for cortical modulation of interoceptive fear of relevance to panic and fear-associated pathologies.

## Acknowledgements

Studies were supported by VA Merit Grants I01-BX001075 (awards 04-12) to RS. KMJM acknowledges support from F32MH117913, R00AA029168 and P50AA022538.

---

**Supplemental Figure 1.** SFO to BNST projection does not regulate acute or delayed CO_2_-associated defensive fear behaviors, startle, neuroendocrine response to stress and dehydration induced water intake. (**A**) Experimental layout schematic and (**B**) Intersectional chemogenetics strategy to target the SFO-BNST circuit using a Cre-expressing retrogradely transported viral construct - pENN.AAV.hSyn.HI.eGFP-Cre.WPRE.SV40 into the BNST and AAV2-hSyn-DIO-hM3D(Gi)-mCherry virus in the SFO to enable Cre-dependent expression of DREADDs in SFO neurons projecting to the BNST. (**C**) Images from the SFO show the expression of Cre-eGFP (green) in Cre expressing SFO to BNST neurons, Cre-dependent Gi-DREADD-mCherry or control mCherry virus expressing red fluorescent protein soma (red) and colocalized cells in merged image. For pre-CO_2_ day context habituation, no significant group difference was observed in freezing (**D**) or rearing (**E**). During 5% CO_2_ inhalation no significant group differences were observed in freezing (**F**) or rearing (**G**). Context re-exposure on day 3 revealed no significant group differences in freezing (**H**) or rearing (**I**). **(J**) CNO administration prior to CO2 inhalation one week earlier had no effect on acquisition in SFO-BNST mice as no group differences were observed following footshocks (**K**) No significant difference in 24 h post-shock conditioned freezing was observed between groups (**L**) Comparison of extinction freezing over 5 days also revealed comparable freezing between groups. (**M**) Mean startle amplitude to a range of acoustic stimuli ranging across 95, 100, 110 and 120[db was not significantly different between groups. Significant effects of decibel but no significant group differences were noted. (**N**) Cumulative water consumption after 24h dehydration stress was comparable between Gi-CNO and control groups as no significant group differences were observed (**O**) Blood corticosterone levels following acute psychogenic restraint stress revealed no significant difference was observed between Gi-CNO, Gi-vehicle and CTRL-CNO groups. Data are expressed as mean ± sem (n= 4-9 mice/group) ns=not significant.

**Supplemental Figure 2.** Prior exposure to CO2 inhalation but not dehydration impacts fear conditioning and extinction. (**A**) Experimental layout shows exposure of separate cohorts to either air/CO_2_ inhalation or 24h water deprivation. One week later mice were exposed to a footshock contextual fear conditioning and extinction paradigm. (**B**) Water-deprived mice showed a significant reduction in body weight and food intake (**C**) supporting that dehydration was a homeostatic threat. One week later, mice with a prior history of acute 24h dehydration elicited comparable freezing to the control group during fear acquisition (**D**), conditioned fear (**E**) and extinction (**F**). In contrast, prior single exposure to CO_2_ a week earlier produced significantly higher freezing in CO_2_ exposed mice during fear acquisition (**J**) conditioned fear (**K**) and extinction (**L**). Data are expressed as mean ± sem (Suppl Fig 2B-F n= 10-12 mice/group; Suppl Fig2G-L n=26-32 mice/group) * p<0.05 **p<0.01 ns=not significant.

**Supplemental Figure 3.** Effects of CO_2_ inhalation in the IL and BNST. **(A)** Layout for experiment on CO_2_ or air inhalation in Fos ^CreERT2.Ai9^ mice. Tamoxifen was injected 1 hour after inhalation. One week later Fos/td-tomato^+ve^ cells were quantified in the BNST. (**B**) Bar graph shows no significant group difference between CO_2_ and air mice within the BNST No significant correlation of Fos/td-tomato^+ve^ cells was observed with freezing (**C**) or rearing (**D)** behaviors during inhalation. (**E**) To determine neuronal response during CO_2_ inhalation we recorded pre and post CO_2_ changes with IL-targeted infusion of a CaMKII promoter driven GCaMP6f and IL-implanted fiberoptic cannula. Analysis for baseline vs CO_2_ revealed a reduction in mean transient amplitude (**F**) and frequency (**G**). Optogenetic activation of SFO CAMKII expressing neurons does not impact somatostatin SST colabelled cfos cells. (H) Image shows co-labelling of SST+/cFos cells in the IL. No difference in cFos + somatostatin (SST) co-labeled neurons was observed between the IL (**I**) or in the PL (**J**) of ChR2 mice vs control mice. Data are expressed as mean ± sem (n= 3-6 mice/group). Photometry data in panels F,G represents within subject recording of 32-50 (mean 44) events per timepoint (n=1) * p<0.05 between pre-CO_2_ and all other times sampled; ns=not significant.

