## Supplementary figures and images for "Projections from subfornical organ to infralimbic cortex modulate carbon dioxide associated fear"

### Supplementary Fig 1

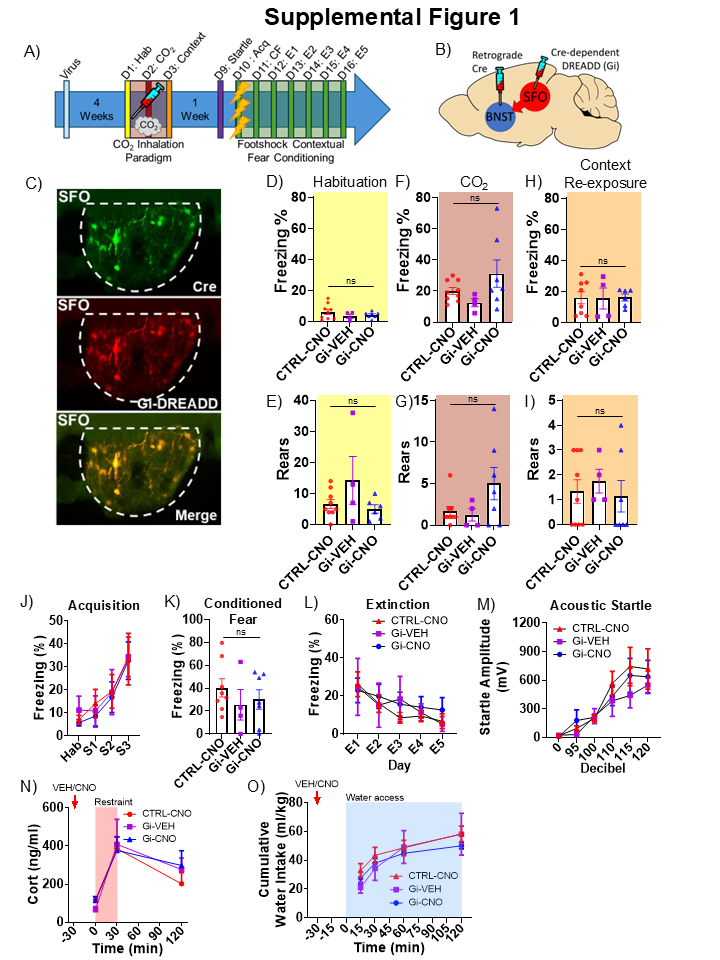

### Supplementary Fig 2

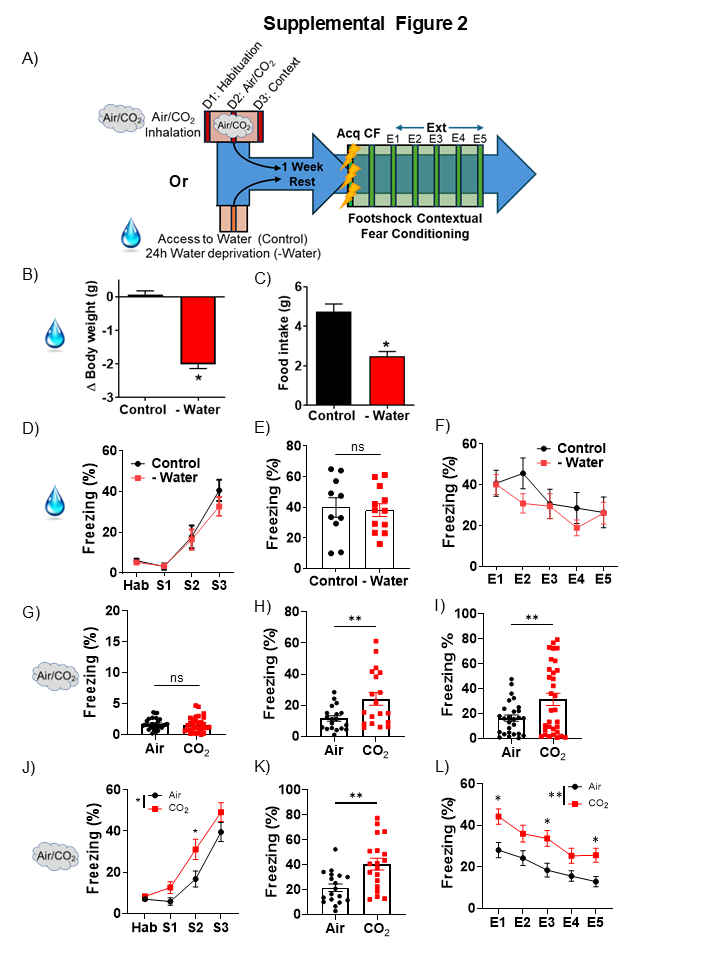

### Supplementary Fig 3

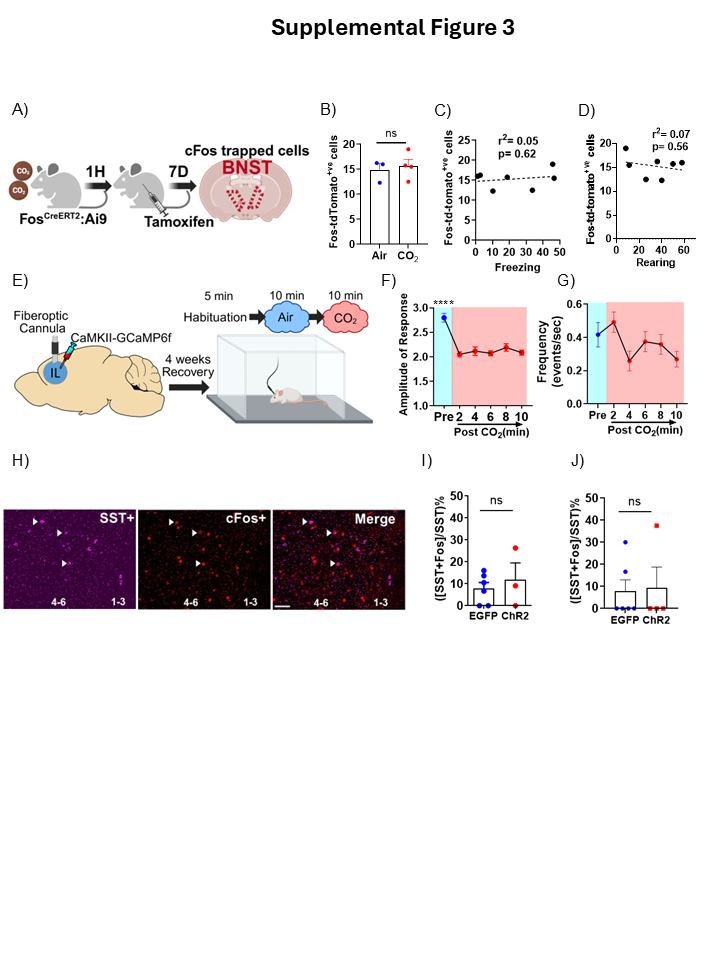
